# Adaptive evolution is substantially impeded by Hill-Robertson interference in Drosophila

**DOI:** 10.1101/021600

**Authors:** David Castellano, Marta Coronado, Jose Campos, Antonio Barbadilla, Adam Eyre-Walker

## Abstract

It is known that rates of mutation and recombination vary across the genome in many species. Here we investigate whether these factors affect the rate at which genes undergo adaptive evolution both individually and in combination and quantify the degree to which Hill-Robertson interference (HRi) impedes the rate of adaptive evolution. To do this we compiled a dataset of 6,141 autosomal protein coding genes from Drosophila, for which we have polymorphism data from *D. melanogaster* and divergence out to *D. yakuba*. We estimated the rate of adaptive evolution using a derivative of the McDonald-Kreitman test that controls for the slightly deleterious mutations. We find that the rate of adaptive amino acid substitution is positively correlated to both the rates of recombination and mutation. We also find that these correlations are robust to controlling for each other, synonymous codon bias and gene functions related to immune response and testes. We estimate that HRi reduces the rate of adaptive evolution by ∼27%. We also show that this fraction depends on a gene’s mutation rate; genes with low mutation rates lose ∼11% of their adaptive substitutions while genes with high mutation rates lose ∼43%. In conclusion, we show that the mutation rate and the rate of recombination, are important modifiers of the rate of adaptive evolution in Drosophila.

## Introduction

It has been shown that there are substantial levels of adaptive protein evolution in many species; for example, in species of Drosophila, rodents, bacteria and some plants it has been estimated that >25% of all amino acid substitutions are consequence of positive adaptive evolution (Bustamante et al. 2002; Smith and Eyre-Walker 2002; Bierne and Eyre-Walker 2003; Sawyer et al. 2003; Charlesworth and Eyre-Walker 2006; Haddrill et al. 2010; Ingvarsson 2010; Slotte et al. 2010; Strasburg et al. 2011). In contrast there are some species, such as humans and many other plants for which rates of adaptive evolution appear to be very low (Chimpanzee Sequencing and Analysis Consortium 2005; Zhang et al. 2005; Boyko et al. 2008; Eyre-Walker and Keightley 2009; Gossmann et al. 2010). The reasons for this variation between species is not fully understood, although effective population size (*Ne*) appears to be important (Gossmann et al. 2012).

The rate of adaptive evolution also appears to vary between genes within a genome. This is expected for several reasons. First, some genes are expected to undergo more adaptive evolution because of their functions; in particular those genes that interact with the environment or which are caught up in arms races are expected to have high rates of adaptive evolution, whereas those genes with highly conserved functions are expected to adapt slowly. Second, genes with high mutation rates are predicted to adapt faster than those with low mutation rates. This is expected whether most adaptation comes from newly arising mutations or from standing genetic variation. This is obvious if adaptation is mutation limited; if an organism is waiting for advantageous mutations to arise, and adaptation can potentially occur in more than one gene, then adaptation is mostly likely to occur in the gene with the highest mutation rate. However, we also expect adaptation to be greater even if advantageous mutations are selected from standing genetic variation, because genes with the highest mutation rates will contribute most to diversity. Third, we expect the rate of adaptive evolution to depend upon the rate of recombination; genes with low rates of recombination will suffer from Hill-Robertson interference (HRi) (Hill and Robertson 1966; Felsenstein 1974) in which selected mutations interfere with each other: a newly arising advantageous mutation may find itself in linkage disequilibrium with deleterious mutations, which will reduce its probability of fixation if it can not recombine away from them, or in competition for fixation with another advantageous mutation at a linked locus on another chromosome in the population. Fourth, we expect an interaction between the rate of recombination and the rate of mutation; HRi should be more prevalent in genes with high mutation rates and low rates of recombination.

Several studies have shown that gene function is important in determining the rate of adaptive evolution: Obbard et al. 2009 have shown that immune system genes have higher rates of adaptive evolution than other genes in Drosophila and Haerty et al. 2007 and Pröschel et al. 2006 have shown that male-biased genes, like testes specific genes, have higher rates of adaptive evolution. It has also been shown that in humans many of the genes that present a signature of positive selection tend to be involved in sensory perception, immune defenses, tumor suppression, apoptosis, and spermatogenesis (Clark et al. 2003; Chimpanzee Sequencing and Analysis Consortium 2005; Nielsen et al. 2005). The role of recombination has also been studied; it has been shown in Drosophila that the rates of adaptation in different regions of the genome vary greatly by differences in the frequency of recombination (Presgraves 2005; Betancourt et al. 2009; Arguello et al. 2010; Mackay et al. 2012; Campos et al. 2014). Surprisingly the role of the mutation rate in the rate of adaptive evolution has not been considered before.

In this work we investigate the effect of the recombination rate and the mutation rate on the rate of adaptive amino acid substitutions (K_a+_) in the genome of *Drosophila melanogaster* and quantify the extent to which HRi impedes the rate of adaptive evolution. Moreover, we are able to estimate how the fraction of lost adaptive amino acid substitutions to HRi depends on gene’s mutation rate.

## Results

To investigate the role of recombination and mutation in determining the rate of adaptive evolution we compiled 6,141 autosomal protein coding genes from Drosophila for which we have polymorphism data from *D. melanogaster* and divergence out to *D. yakuba.* For most of our analyses we use polymorphism data from the *Drosophila melanogaster* Genetic Reference Panel (DGRP) which was sampled from Raleigh, North Carolina (Mackay et al. 2012). However, in some analyses we compare our results to those obtained using the flies sampled from Gikongoro, Rwanda (DPGP2, Pool et al. 2012). To estimate the rate of adaptive evolution we use the DFE-alpha method (Eyre-Walker and Keightley 2009), a derivative of the McDonald-Kreitman test (McDonald and Kreitman 1991) which corrects for slightly deleterious mutations. In this method it is assumed that mutations at one set of sites (in this analysis synonymous sites) are neutral and that selection acts upon the mutations at another set of sites (non-synonymous sites). The site frequency spectra (SFS) of synonymous and non-synonymous polymorphisms are used to infer the distribution of fitness effects of neutral and deleterious mutations at the non-synonymous sites and this information is used, in conjunction with the level of synonymous divergence, to predict how many neutral and nearly neutral non-synonymous substitutions are expected. If the observed divergence at non-synonymous sites exceeds this expectation, adaptive evolution is inferred and quantified. The rate of adaptive evolution is typically estimated using one of three statistics: K_a+_, the rate of adaptive amino acid substitution, ω_A_, the rate of adaptive evolution relative to the mutation rate and α, the proportion of substitutions that are adaptive. The α statistic conflates the rates of adaptive and non-adaptive substitution and hence is not useful for our purposes here, and ω_A_ is not useful for studying the effects of mutation on the rate of adaptive evolution because it controls for the factor being investigated, hence we have investigated how K_a+_ depends upon the rate of recombination and mutation. However, in terms of recombination, we get qualitatively similar results whether we use K_a+_ or ω_A_.

### Recombination and Adaptation

We first studied the relationship between recombination rate and K_a+_. To estimate the rate of adaptive evolution it is necessary to combine data from several genes because estimates tend to be error prone and sometimes undefined for individual genes. We therefore grouped genes into 45 bins of 136 genes each based on their rates of recombination. The results are shown in fig. 1A and supplementary Table 1 A. There is a highly significant positive relationship between the rate of adaptation and recombination rate (Spearman’s rank correlation coefficient *ρ* = 0.64, p < 0.001). However, for values beyond ∼2 cM/Mb the relationship between recombination and adaptation reaches an asymptotic value. In order to test whether a curvilinear relationship fits the data better than a linear model we fit the function *y* = *a* + *b* · *e*^−*cx*^ to our data and compared it to the fit of a linear model. Table 1 shows the the inferred parameters, the R^2^ and AIC values for the two models. In terms of both AIC and R^2^ the curvilinear model is favoured.

**Table 1.**
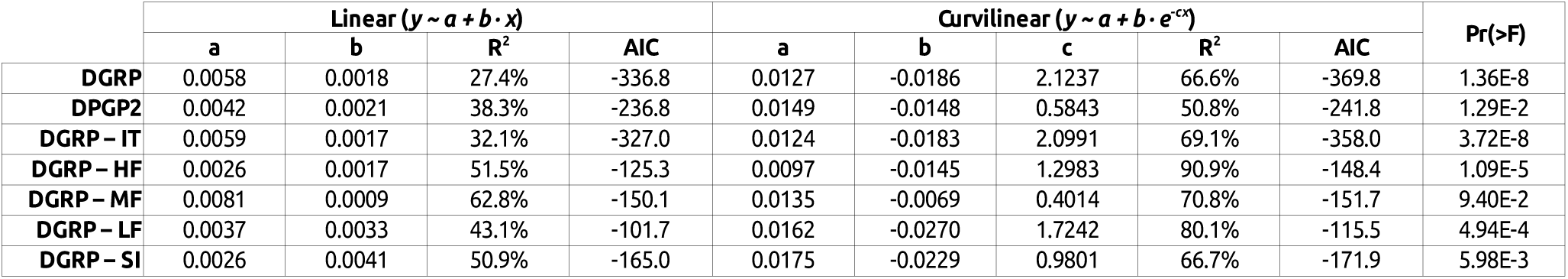
Linear and curvilinear fit inferred parameters, *R*^2^ and AIC for seven data sets where y = K_a+_ and x = cM/Mb. The last column is the p-value for a F-test comparing both models. The DGRP data set is in the first row, the DPGP2 data is in the second row, the DGRP data excluding immune response and testes related genes is in the third row, the DGRP data for high, medium and low *Fop* genes are in the fourth, fifth and sixths rows, respectively, and finally the DGRP data using short intron sites (<66, bases 8-30) as neutral standard is in the last row.

| | Linear ($y \sim a + b \cdot x$) | | | | Curvilinear ($y \sim a + b \cdot e^{-cx}$) | | | | | Pr(>F) |
| --- | --- | --- | --- | --- | --- | --- | --- | --- | --- | --- |
|  | a | b | R<sup>2</sup> | AIC | a | b | c | R<sup>2</sup> | AIC |  |
| **DGRP** | 0.0058 | 0.0018 | 27.4% | -336.8 | 0.0127 | -0.0186 | 2.1237 | 66.6% | -369.8 | 1.36E-8 |
| **DPGP2** | 0.0042 | 0.0021 | 38.3% | -236.8 | 0.0149 | -0.0148 | 0.5843 | 50.8% | -241.8 | 1.29E-2 |
| **DGRP – IT** | 0.0059 | 0.0017 | 32.1% | -327.0 | 0.0124 | -0.0183 | 2.0991 | 69.1% | -358.0 | 3.72E-8 |
| **DGRP – HF** | 0.0026 | 0.0017 | 51.5% | -125.3 | 0.0097 | -0.0145 | 1.2983 | 90.9% | -148.4 | 1.09E-5 |
| **DGRP – MF** | 0.0081 | 0.0009 | 62.8% | -150.1 | 0.0135 | -0.0069 | 0.4014 | 70.8% | -151.7 | 9.40E-2 |
| **DGRP – LF** | 0.0037 | 0.0033 | 43.1% | -101.7 | 0.0162 | -0.0270 | 1.7242 | 80.1% | -115.5 | 4.94E-4 |
| **DGRP – SI** | 0.0026 | 0.0041 | 50.9% | -165.0 | 0.0175 | -0.0229 | 0.9801 | 66.7% | -171.9 | 5.98E-3 |

**Fig. 1.**
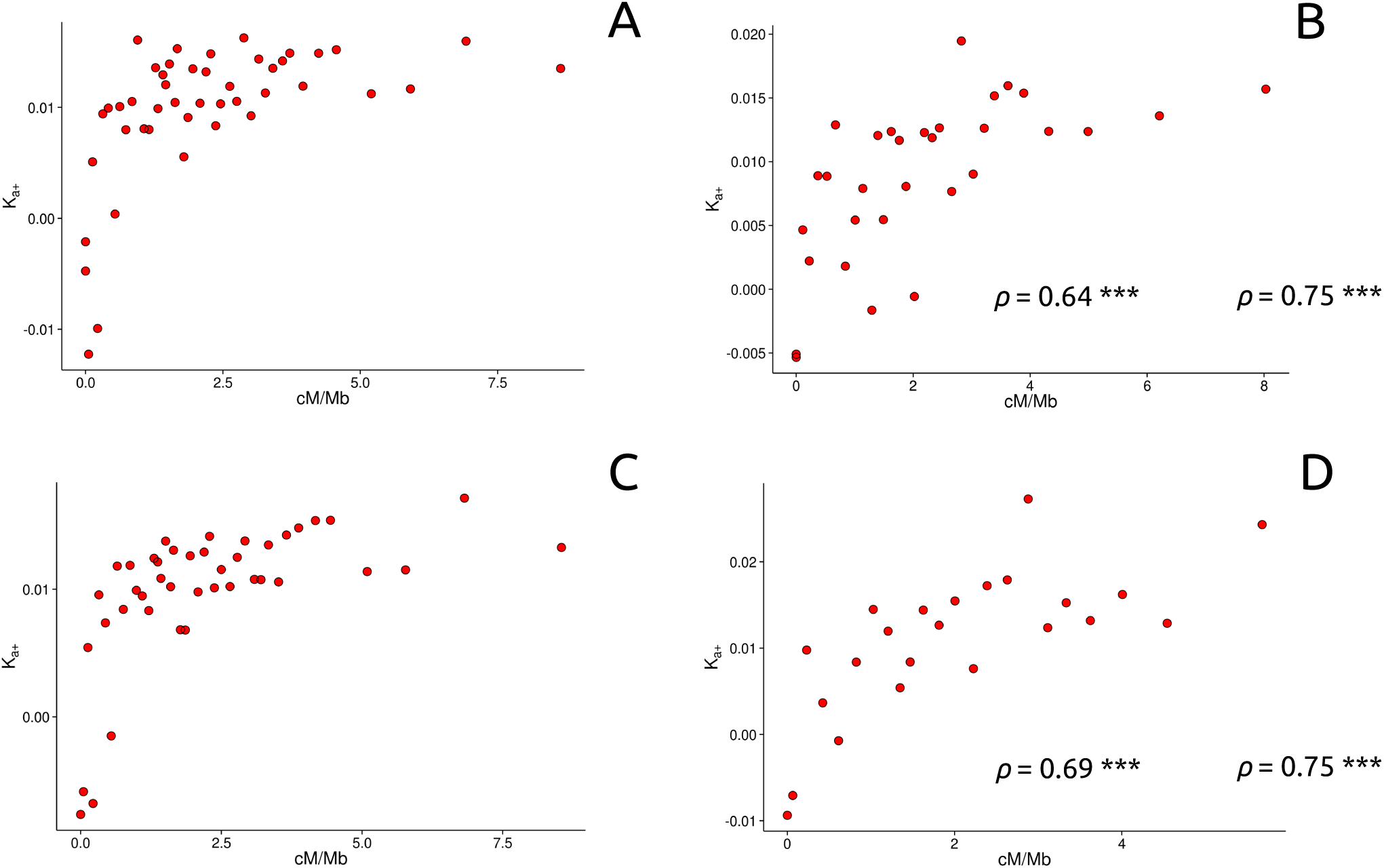
Relations between K_a+_ in the *y* axis and the rate of recombination (cM/Mb) in the *x* axis: (A) using DGRP data, North Carolina population, (B) using DPGP2 data, Rwanda population, (C) excluding immune response and testes related genes and (D) using as neutral reference short intron sites (< 66 nt, bases from 8 to 30). Each data point has been estimated binning 136 genes according to their recombination rate levels. The recombination rate ranks can be consulted in the supplementary table 1. *ρ*: Spearman’s rank correlation coefficient, with significance denoted by asterisks (***<0.001; **<0.01; *<0.05).

Our results are in contrast to those of Campos et al. 2014 who found that ω _A_ was linearly related to the rate of recombination. The difference between the two analyses could be due to the fact that we have used a different measure of adaptive evolution, the polymorphism data come from different populations or to differences in the way in which the rate of recombination was estimated. The difference between the two studies is not due to the measure of adaptive evolution used; we observe a curvilinear relationship using both K_a+_ and ω_A_ (see supplementary fig. 1). Campos et al. 2014 used two different estimates of recombination rate: one based on low resolution visible markers (Fiston-Lavier et al. 2010), the other one on the high resolution recombination map obtained by Comeron et al. 2012 using SNP markers. For both datasets Campos et al. 2014 observe a linear relationship. However, instead of taking point estimates of the recombination rate from the Comeron et al.’s high resolution map as we have done, Campos et al. 2014 fitted a LOESS regression to the data which smoothes out the original high resolution recombination map. We have repeated the correlation analysis of Campos et al.’ using their polymorphism (from Gikongoro, Rwanda [DPGP2, Pool et al. 2012]) and divergence genomic data together with the original unsmoothed high resolution recombination map. In contrast to the linear relationship they originally reported we found the same highly significant curvilinear pattern that we observed using the DGRP polymorphism data (see fig. 1B, Table 1 and supplementary Table 1 B) (Spearman’s rank correlation coefficient *ρ* = 0.75, p < 0.001). Thus, the linear relationship between the rate of adaptive evolution and the rate of recombination observed by Campos et al. seems to be a consequence of smoothing the recombination rate estimates rather than differences in the populations from which the polymorphism data were derived. For this reason, all subsequent analyses presented here are based on the DGRP data from Raleigh, North Carolina (Mackay et al. 2012), since these data have a greater coverage and number of sampled chromosomes.

The positive correlation between recombination rate and K_a+_ could be due to a number of potential biases in the dataset. For example, if some classes of genes with high rates of adaptation are preferentially located in regions with high rates of recombination, the observed positive correlation would not necessarily be a consequence of adaptive evolution being affected by recombination. There is evidence that immune system (Obbard et al. 2009) and male-biased or testes specific (Pröschel et al. 2006; Haerty et al. 2007) genes undergo higher rates of adaptive evolution than other genes. We confirm this result taking into account the influence of slightly deleterious mutations, which the previous analyses did not. We find that immune and testes-specific genes together exhibit adaptive rates 1.37x faster than other genes (the difference between immune/testes specific genes and other “control” genes is significant as judged by a permutation test – p = 0.017). However, if we remove immune and testes specific genes we still observe a highly significant curvilinear correlation between recombination rate and K_a+_ (see fig. 1C, Table 1 and supplementary Table 1 C) (Spearman’s rank correlation coefficient *ρ* = 0.69, p < 0.001).

In estimating the rate of adaptive evolution we have assumed that synonymous mutations are neutral, however selection is known to act upon synonymous sites in Drosophila (reviewed by Hershberg and Petrov 2008). In many cases, this is thought to be a result of selection favoring codons that can be translated more rapidly or accurately (Shields et al. 1988; Akashi 1994, 1995; Carlini and Stephan 2003; Stoletzki and Eyre-Walker 2006). Additionally, synonymous sites may be under selection to maintain (or avoid) splicing enhancers (Parmley et al. 2006), messenger RNA secondary structures (Parsch et al. 1997; Baines et al. 2004; Stoletzki 2008) or particular short sequence motifs (Antezana and Kreitman 1999). Lawrie et al. 2013 have shown that ∼22% of all 4-fold synonymous sites in *D. melanogaster* are under strong purifying selection, although the specific functional mechanism underlying this strong constraint is complex. Consistent with this weak selection favouring codons that have to be translated more rapidly or accurately we confirm previous results that K_4_ is significantly correlated to a measure of codon usage bias, *Fop* (the frequency of optimal codons) (Spearman’s rank correlation coefficient ρ = -0.4, p < 0.001) (Sharp and Li 1987, 1989; Moriyama and Hartl 1993; Bierne and Eyre-Walker 2003, 2006). However, we expect that any sort of weak selection on synonymous mutations to generate a positive correlation between K_a+_ and recombination rate. This is because we expect that genes located in regions of high recombination, where selection on synonymous sites is more efficient (Kliman and Hey 1993; Haddrill et al. 2007; Campos et al. 2012), will tend to have a higher estimate of K_a+_ because weak negative selection on synonymous mutations inflates the number of synonymous polymorphisms relative to the number synonymous substitutions. Therefore, to investigate whether selection on synonymous codon usage affects our adaptation estimates we divided our genes into 3 roughly equal groups according to their *Fop* value, and within each of these *Fop* groups we divided the data into 15 groups of genes according to their recombination rate (details of the groups can be found in the online supplementary material). We observe the same highly significant curvilinear relationship within each of the 3 *Fop* categories (see supplementary fig. 2, Table 1 and supplementary Table 3) (Spearman’s rank correlation coefficient for high *Fop* genes *ρ* = 0.87, p < 0.001; medium *Fop* genes *ρ* = 0.78, p < 0.001; low *Fop* genes *ρ* = 0.76, p < 0.001). We also repeated our analysis using a smaller dataset of 3,369 genes where we can use polymorphisms and substitutions in short introns (<66 bp) as the neutral standard. This dataset is smaller because not all genes fulfil the intron quality and length criteria (see *Materials and Methods*). The same curvilinear pattern is observed (see fig. 1D, Table 1 and supplementary Table 1 D) and the strength of the correlation is equivalent to that found with 4-fold sites (Spearman’s rank correlation coefficient *ρ* = 0.75, p < 0.001). Hence, selection on codon usage does not seem to be responsible for the shape or the strength of the relationship between the rates of adaptive evolution and recombination.

**Fig. 2.**
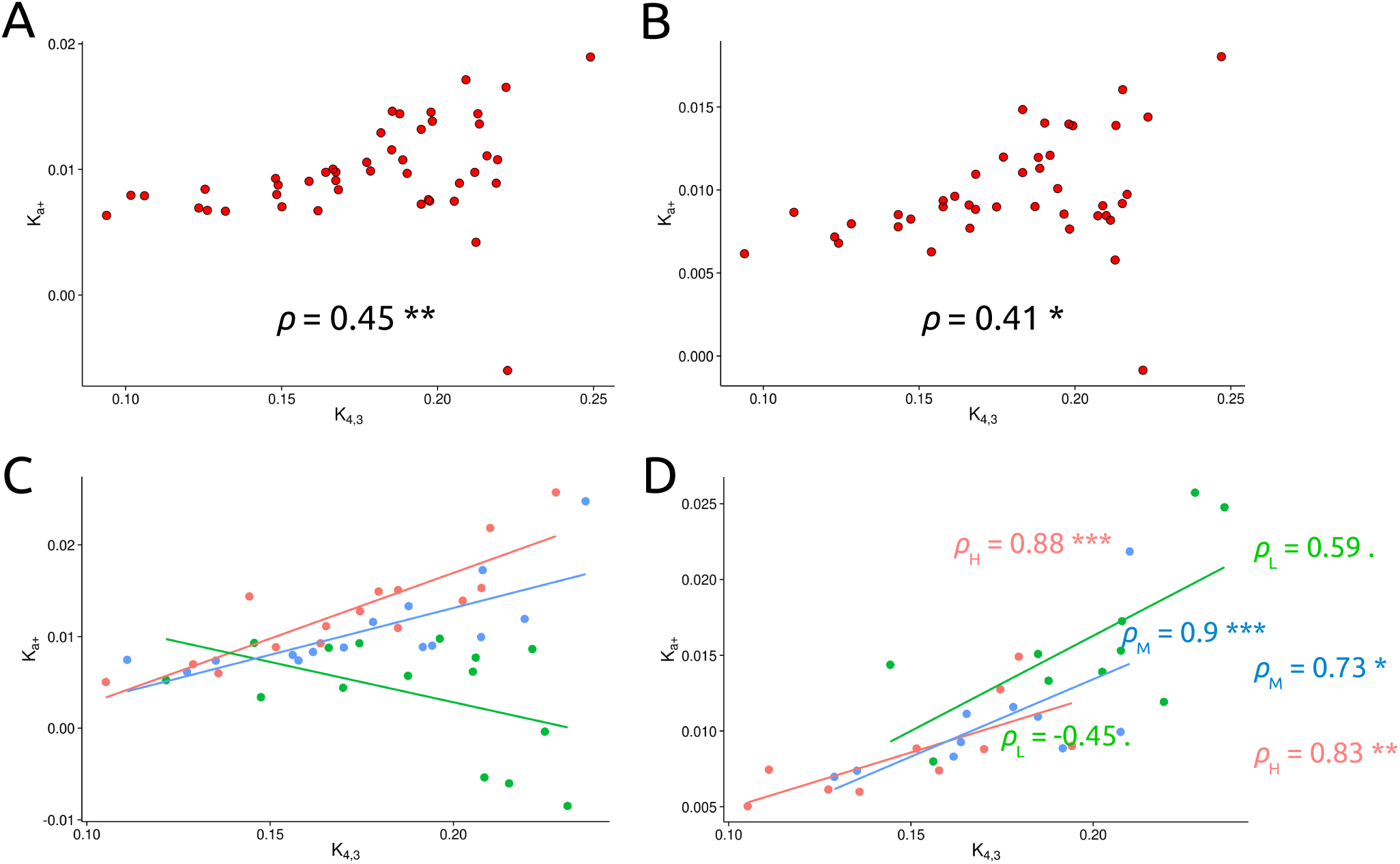
Relationship between K_a+_ in the *y* axis and an estimate of the mutation rate (K_4,3_) in the *x* axis: (A) using DGRP data, North Carolina population, (B) excluding immune response and testes related genes, (C) splitting the dataset into three recombination groups and (D) splitting the dataset into three *Fop* groups after removing low recombination rate genes (< 1.32 cM/Mb). Genes belonging to the high (H) group are in red, medium (M) genes are in blue and low (L) genes are in green. Each data point has been estimated binning 136 genes according to their mutation rate (K_4,1_). The mutation rate ranks can be consulted in the supplementary table 2. *ρ*: Spearman’s rank correlation coefficient, with significance denoted by asterisks (***<0.001; **<0.01; *<0.05;. 0.1-0.05). The lines are least-squares regressions but should be regarded only as indicative, in view of the binning of the data.

### Mutation and Adaptation

To investigate whether the rate of adaptive evolution is correlated to the mutation rate is not straightforward since we need to use the rate of synonymous substitution to estimate both the mutation rate and the rate of adaptive evolution. This lack of statistical independence between estimates will tend to generate a negative correlation between K_a+_ and K_4_ just through sampling error. To avoid problems of non-independence we split our synonymous substitution estimate, K_4_, into 3 independent variables by sampling from a hypergeometric distribution (see *Materials and Methods: Hypergeometric Sampling*); we used K_4,1_ to rank genes and assign genes to bins, K_4,2_ to estimate the rate of adaptive evolution, and K_4,3_ as an estimate of mutation rate for this bin.

The data were divided, as with the recombination rate analyses, into 45 mutation bins of 136 genes each, but this time the data were divided according to their K_4,1_ value. Doing this we found a highly significant positive correlation between K_a+_ and K_4,3_ (Spearman’s rank correlation coefficient *ρ* = 0.45, p < 0.001, see fig. 2A and supplementary Table 2 A). As with the correlation between K_a+_ and the rate of recombination, this correlation could be spurious due to several sources of bias. The correlation is still highly significant even if we exclude testes and immune system related genes (Spearman’s rank correlation coefficient *ρ* = 0.41, p < 0.01, see fig. 2B and supplementary Table 2 B), suggesting that the correlation between K_a+_ and K_4_ is not a consequence of the non-random distribution of this kind of genes relative to the mutation rate. Natural selection on codon usage is expected to weaken rather than generate an artefactual positive correlation between K_a+_ and K_4,3_, because selection on codon usage should reduce the rate of synonymous substitutions more than the level of synonymous polymorphism. To investigate whether selection on codon usage has an effect on the relationship between K_a+_ and K_4,3_ we divided the data set into 3 recombination rate levels and 3 *Fop* levels, and within each recombination rate and *Fop* class we grouped the genes into 5 groups according to their mutation rate (this yielded 45 bins of 136 genes each, see supplementary Table 2 C). We separate the data according to their recombination rate because it affects both the rate of adaptive evolution as well as the efficiency of selection on codon usage. The correlation between K_a+_ and K_4,3_, for each recombination rate and *Fop* category is shown in figure 2C and 2 D, respectively. The graphs suggest that selection on codon bias makes little difference to the correlation between K_a+_ and K_4_, but that the relationship is strongly affected by the rate of recombination; this is not surprising because we have shown above that genes with low rates of recombination undergo very little adaptive evolution (see fig. 1) and therefore not likely to be influenced by the rate of mutation. To investigate this more formally we performed an analysis of covariance (ANCOVA), grouping genes by their *Fop* and recombination rate levels. In ANCOVA, a set of parallel lines are fitted to the data, one for each group. This enables a test of whether the common slope of these lines is significantly different from zero, and one can also investigate whether the groups differ in the dependent variable for a given value of the independent variable by testing whether the lines have different intercepts. If we consider *Fop* and recombination rate as fixed factors we find no significant correlation between K_a+_ and K_4,3_ (ANCOVA p = 0.16). However, we find evidence that the slopes (ANCOVA p < 0.001) and intercepts (ANCOVA p < 0.001) differ between recombination rate categories, but there is no evidence that either the slope or intercept differs between *Fop* categories. If genes with low recombination rates (from 0 to 1.32 cM/Mb) are excluded, a very strong positive correlation between K_a+_ and K_4,3_ is found for the rest of the dataset (see supplementary fig. 3 and supplementary Table 2 C) (Spearman’s rank correlation coefficient *ρ* = 0.82 and p < 0.001). There is no evidence within this dataset that the slope or intercept differ according to rate of recombination or the level of codon bias. As an alternative approach to controlling the effect of selection on codon usage on our estimates of the mutation rate we regressed K_4,3_ against the rate of recombination and *Fop* and used the residuals as a measure of the mutation rate. We find a strong positive correlation between K_a+_ and the residuals (see supplementary fig. 4 and supplementary Table 2 A) (Spearman’s rank correlation coefficient ρ = 0.42 and p < 0.01), suggesting that the correlation between K_a+_ and K_4_ is not a result of weak selection on 4-fold sites.

### The Proportion of Adaptive Substitutions Lost to HRi

Our results show that the rate of adaptive evolution is significantly impeded in low recombining regions of the Drosophila genome. But how many adaptive substitutions are lost because of HRi? And how does the mutation rate affect the intensity of the HRi? To answer these questions we fit a LOESS curve to the relationship between K_a+_ and recombination rate, which clearly approaches an asymptote above 2cM/MB. The asymptote > 2cM/Mb can be interpreted as the rate of adaptive evolution that would occur if there was no HRi. The LOESS curve decreases below the asymptotic value as the rate of recombination decreases, and the difference between the asymptote and the LOESS curve can be interpreted as the number of adaptive substitutions that are lost due to HRi. Using this approach we estimate, after weighting by the number of sites involved that 27.2% (95% CIs obtained by bootstrapping by gene [20.6%, 33.8%]) of all adaptive amino acid substitutions that would be fixed in an effectively free recombining genome are lost because of HRi. Some of the estimates of K_a+_ inferred from the LOESS curve are negative; however, even our estimate of the proportion of adaptive substitutions lost to HRi is largely unchanged even if we set these to zero: 27.1% (95% CIs [20.6%, 33.2%]).

HRi is expected to be more prevalent in regions of the genome with higher rates of mutation, because this will increase the chance that a selected mutation will be segregating with other mutations subject to selection. To investigate whether this is the case in Drosophila we repeated the analysis above splitting the dataset into two subsets according to a gene’s inferred mutation rate. To do this we split K_4_ into two independent variates by sampling from a hypergeometric distribution. K_4,1_ was used to divide the genes into different mutation rate categories, while K_4,2_ was used to calculate K_a+_. In this way we ensured that the estimates of adaptive evolution were not influenced by the way in which the data was divided. We find that the proportion of adaptive substitutions differs significantly between the 25% of genes with lowest and the 25% with highest mutation rate estimates (p-value from a bootstrap analysis = 0.002): the bottom 25% of genes are estimated to have lost 10.9% (95% CIs [-0.3%, 24.4%]) of their adaptive amino acid substitutions to HRi whereas the top 25% have lost 42.5% (95% CIs [30.7%, 55%]). Alternatively, to control for the effect of selection on synonymous sites we regressed K_4,1_ against the rate of recombination and *Fop* and used the residuals as a measure of the mutation rate to rank and group genes. The bottom 25% of genes have lost 18.1% (95% CIs [4.8%, 31.2%]) of their adaptive amino acid substitutions whereas the top 25% have lost 41.3% (95% CIs [27.9%, 55.5%]). Again, the proportion of adaptive substitutions differs significantly between both kind of genes (bootstrap p-value = 0.028). In any case, the average fraction of lost adaptive substitutions for the low (and high) mutation rate genes are not significantly different between both estimates of the mutation rate. This result indicates that gene’s mutation rate affects how influential HRi is.

## Discussion

We have shown that the rate of adaptive protein evolution is positively correlated to both the rate of recombination and the mutation rate in *D. melanogaster*. We have shown that these correlations are not due to an enrichment of immune response and testes related genes in regions of high recombination or mutation, or due to selection on synonymous sites. Instead it seems likely that the rate of adaptive evolution is correlated to the rate of recombination because of Hill-Robertson interference (HRi) and that it is correlated to the rate of mutation because genes with higher rates of mutation are more likely to produce the genetic variation needed for adaptation. This work quantifies for the first time the global impact of HRi on a given genome. We estimate that approximately 27% of all adaptive mutations, which would go to fixation if there was free recombination, are lost due to HRi. We show that this effect depends upon the mutation rate with genes with high mutation rates losing a greater proportion of their adaptive substitutions to HRi than genes with low mutation rates.

The recombination rate data we have used only includes cross-overs (CO) and excludes gene conversion (GC) events. This is because GC is expected to be a much less important force reducing HRi than CO. Although GC events occur ∼5-times more frequently than COs (Comeron et al. 2012), the GC tract lengths are quite short at about 500bp (Comeron et al. 2012) and hence lead to relatively little recombination. The fact that GC is largely ineffective in reducing HRi can be inferred from the presence of HRi in regions of the genome with very low rates of CO, because even these regions have moderate levels of GC – the frequency of GC varies little across the Drosophila genome (Comeron et al. 2012).

An open question is to what extent HRi affects rates of adaptive evolution in other species. The strength of HRi depends on the rate of mutation, the distribution of fitness effects (DFE) and the rate of recombination; the greater the density of selected mutations per map unit, and the more strongly selected they are, the greater the effect of HRi on weakly selected mutations. Is HRi likely to be an important force in a species like humans? Humans are estimated to have a genomic rate of harmful mutation of 2.1 (Lesecque et al. 2012) that is approximately twice that in Drosophila at 1.2 (Haag-Liautard et al. 2007), and although, the human genome is approximately 20x greater in size than the Drosophila genome, linkage disequilibrium declines approximately 500x more slowly in humans than Drosophila. Taken together these results suggest that HRi, at least from deleterious mutations, might be more important in humans than Drosophila. However, this needs to be confirmed by analysis, and this is difficult because humans appear to have undergone relatively little adaptive evolution (Boyko et al. 2008; Eyre-Walker and Keightley 2009; Gossmann et al. 2012) and this makes analysing the factors that affect the rate of adaptive evolution difficult. The potentially higher level of HRi in humans may explain in part why our species appears to have undergone relatively little adaptive evolution compared to Drosophila (Gossmann et al. 2012). However, the effect of HRi will depend upon the distributions of fitness effects and this is something we have limited information about in both of these species. It will be of great interest to do similar analyses to those performed here in other species.

The loss of adaptive substitutions to HRi can potentially tell us something important about the strength of selection acting on some advantageous mutations, since weakly selected mutations are those that are most likely to be affected by HRi (McVean and Charlesworth 2000; Comeron and Kreitman 2002; Comeron et al. 2008). This will require further analysis and population genetic modelling, but in combination with other sources of information, for example, the dip in diversity around non-synonymous substitutions (Sella et al. 2009), the site frequency spectrum (Schneider et al. 2011) the high frequency variants that are left by selective sweeps (Fay and Wu 2000) it may be possible to infer much more about the DFE of advantageous mutations than previously thought.

The fact that so many adaptive substitutions are lost to HRi begs the question why Drosophila does not have a higher rate of recombination, particularly in areas where there is little or no recombination in its genome. This may be because selection on modifiers of the recombination rate is weak; a modifier that elevates the rate of recombination may allow advantageous mutations to spread more easily, but by its very nature it will tend to disassociate itself from the advantageous mutations that it helps spread. It therefore gets little or no benefit from the positive effects it causes.

## Conclusions

This work has shown how two genomic forces, the rate of recombination and mutation rate, affect the rate of adaptation within the *Drosophila melanogaster* genome. We find that the rate of adaptive amino acid substitution is positively correlated to both recombination rate and an estimate of the mutation rate. We also find that this correlation is robust to controlling for each other, synonymous codon bias and gene functions related to immune response and testes. Finally we estimate that on average ∼27% of all advantageous substitutions have been lost because of HRi and that this quantity depends on gene’s mutation rate; ∼11% of adaptive substitutions have been lost for low mutation genes and ∼43% for high mutation genes. Hence, we have shown evidences that both recombination and mutation are important modifiers of the rate of adaptive evolution in Drosophila.

## Materials and Methods

### Population Genomic Data, Polymorphism and Divergence Estimates

This study was carried out on the four large autosomes (2L, 2R, 3L and 3R) of *D. melanogaster* using release 5 of the Berkeley Drosophila Genome Project (BDGP 5, http://www.fruitfly.org/sequence/release5genomic.shtml) as the reference genome.

#### North American Population

The population genomic data comes from Raleigh, North Carolina. The details of their provenance and breeding are in Mackay et al. 2012, Freeze 1.0 *Drosophila melanogaster* Genetic Reference Panel (DGRP) project. Sites with residual heterozygosity and low quality values were excluded from the analyses. The method for jointly estimating the distribution of fitness effects on new mutations (DFE) and the rate of adaptive substitution requires all sites to have been sampled in the same number of chromosomes (DFE-alpha method, Eyre-Walker and Keightley 2009; see below) and since some sites were not successfully sampled in all samples we reduced the original data set to 128 isogenic lines by randomly sampling the polymorphisms at each site without replacement. To estimate divergence out to *D. yakuba* we sampled randomly one *D. melanogaster* single chromosome.

Coding exon and short intron (≤65 bp) annotations from *D. melanogaster* were retrieved from FlyBase (release 5.50, http://flybase.org/, last accessed March 2013). Genes 1:1 orthologs across *D. yakuba* – *D. melanogaster* were obtained from FlyBase (http://flybase.org/). We used *D. yakuba* as outgroup species since there is less chance of ancestral polymorphism contributing to divergence, avoiding in this way the effect of low divergence affecting the estimates of adaptive evolution (Keightley and Eyre-Walker 2012). We obtained a multiple genome alignment between the DGRP isogenic lines (Mackay et al. 2012) and the *D. yakuba* genome (Clark et al. 2007) using the BDGP 5 coordinates. This alignment is publicly available at http://popdrowser.uab.cat/ (Ràmia et al. 2012). For each gene we took all non-overlapping coding exons, independently of their inclusion levels. When two exons overlapped, the largest was chosen for subsequent analyses. Only exons without frameshifts, gaps or early stop codons were retained. In this way we tried to avoid potential alignment errors will inflate our mutation and adaptation rate estimates and create an artefactual positive correlation between them. Our final data set fulfilling all these criteria had 6,141 coding genes.

Exonic sequences were trimmed in order to contain only full codons. We defined our sites “physically”, so we estimated the rates of substitution at sites of different degeneracy separately. Only zero-fold and 4-fold degenerate sites in exon core codons (as described by Warnecke and Hurst 2007) were used. To estimate the rate of synonymous substitutions, we restricted our analysis to those triplets coding the same amino acid in the two species (*D. melanogaster* – *D. yakuba*). In restricting our analysis to codons not exhibiting nonsynonymous differences we assume that the codon has undergone no amino acid substitution — this avoids having to compute the different pathways between two codons, which differ by more than one change and it is a reasonable assumption given the level of amino acid divergence. For 4-fold degenerate sites we used the method of Tamura (Tamura 1992) to correct for multiple hits; this method allows for unequal GC content and ts/tv bias. Jukes and Cantor substitution method was used to correct for multiple hits at zero-fold sites (Jukes and Cantor 1969). We calculated the number of substitutions and the folded site frequency spectrum (SFS) for 4-fold degenerate sites and zero-fold degenerate sites, using an ad hoc Perl Script.

Following Halligan and Keightley 2006, in this study we used positions 8–30 of introns ≤65 bp in length as an alternative neutral reference for some analyses. For intron sequences, the invariant GT and AG dinucleotides at the 5′ and 3′ splice junctions, respectively, were excluded before calculating divergence. Only genes with at least two short introns and with less than 10% of gaps in the aligned sequences were kept. 3,369 orthologous genes passed the intron quality criteria in our final data set. We used an ad hoc Perl Script to estimate the number of short intron substitutions and to compute the folded SFS. Multiple hits were corrected using Jukes and Cantor method (Jukes and Cantor 1969).

#### African Population

We also used population genomic data from an African population. This comes from Gikongoro, Rwanda (DPGP2, Pool et al. 2012). The details of the assembly and data filtering can be found in Campos et al., 2014. The number of synonymous and nonsynonymous sites and substitutions (computed by Comeron 1995 method, which defines a site as a “mutational opportunity”) and the SFS for 7,231 autosomal coding genes were estimated by Campos et al. 2012 and details are provided there. We study only those genes shared by both data sets (DGRP and DPGP2) taking into account the differences in gene annotation versions. This resulted in a dataset of 4,283 autosomal genes coming from this dataset.

### Codon Bias Estimates and Recombination Landscape

We used the CodonW software (http://codonw.sourceforge.net/ by Peden 1999) to estimate the frequency of optimal codons, *Fop*. A higher *Fop* value suggest a higher efficacy of selection for codon usage, and vice versa.

Recombination rates were taken from Comeron et al. 2012 (www.recobinome.com). They estimated the rate of crossovers in 100 kb non-overlapping windows in cM/Mb units. The rate of crossing-over for a gene was the rate in the 100kb that overlapped the mid-point of the gene. Unlike Campos et al. 2014, we did not apply LOESS regression to smooth out the recombination landscape, since we were interested in the fine-scale effects of recombination on the *D. melanogaster* genome.

### Testes, Immune Genes and Permutation Test

If immune and male-biased or testes specific genes tend to have higher rates of recombination (and mutation) then the correlations to the rate of adaptive evolution would not necessarily be a consequence of adaptive evolution being affected by recombination (or mutation) rate. Thus, gene Ontology (GO) terms for 6,141 genes were downloaded from Fruitfly release 78 using the R package biomaRt (Durinck et al. 2005). A list of GO terms related to immune response and testes was constructed using the EBI’s GO tool QuickGO (Binns et al. 2009). When a given gene was associated to a GO term from this list it was labelled as “Immune&Testes genes”, the rest of genes were labelled as “Control genes”. The list of immune response and testes related GO terms and the lists of genes in each group can be consulted in the supplementary Table 4. A permutation test was applied to assess whether K_a+_ are significantly higher as it has been reported before (Pröschel et al. 2006; Haerty et al. 2007; Obbard et al. 2009) for immune response and testes related genes relative to the rest of control genes. We shuffled without replacement 1,000 times the complete list of genes by means of ad hoc Bash and Perl Scripting. Then, we estimated K_a+_ using the DFE-alpha software (Eyre-Walker and Keightley 2009, see below) for each randomized group. Thus, we got the expected null distribution for the differences between “Control genes” minus the “Immune&Testes genes” for the statistic K_a+_. Finally, the one-tailed p-value was obtained by counting the number of replicates below the observed difference divided by the total number of replicates (1000). The expected null distributions and the observed differences can be consulted in the supplementary figure 5.

### Gene Bins and Adaptation Estimates

To estimate the rate of adaptive evolution it is necessary to combine data from several genes because estimates from a single gene are noisy and often undefined because of the lack of segregating (or divergent) sites for some site classes. We therefore grouped genes into bins according to their rate of recombination, mutation rate and/or *Fop*. The rank of values for all these bins can be consulted in the supplementary material.

It is essential to have a selection-free reference sequence that can be used as a baseline for determining the the rate of adaptive substitution acting on a particular target sequence (in our case zero-fold degenerate sites). In this study we used the exon core 4-fold degenerate sites as the main proxy for the neutral mutation rate. For some cross-validation analyses short intron sites were also used. DFE-alpha (Eyre-Walker and Keightley 2009) models the DFE at functional sites by a gamma distribution, specified by the mean strength of selection, γ = −Nes, and a shape parameter β, allowing the distribution to take on a variety of shapes ranging from leptokurtic to platykurtic. DFE-alpha can model a single, instantaneous change in population size from an ancestral size N_1_ to a present-day size N_2_ having occurred t_2_ generations ago. Provided the SFS at both neutral and functional sites and the respective levels of divergence, DFE-alpha infers γ, β, N2/N1, t_2_, and α at functional sites. From these estimates K_a+_ can be easily estimated with the expression: K_a+_ = α x K_a_. We ran DFE-alpha for each bin independently using the local version provided at: http://www.homepages.ed.ac.uk/pkeightl//software. DFE-alpha was run in the folded SFS mode since the results are more robust.

### Hypergeometric Sampling

To analyse the role of mutation on adaptation we correlated 4-fold divergence (K_4_) to the rate of adaptive zero-fold substitutions (K_a+_). A limitation here is that K_4_ and K_a+_ are not independent, since the estimation of K_a+_ depends on K_4_, and then we expect *K*_*a*+_ and *K*_*4*_ to be negatively correlated just through sampling error. To overcome this problem we split our mutation rate estimate (K_4_) into 3 independent variables (similar to the splitting done in (Smith and Eyre-Walker 2002; Piganeau and Eyre-Walker 2009; Stoletzki and Eyre-Walker 2011; Gossmann et al. 2012). This was done by generating a random multivariate hypergeometric variable as follows:

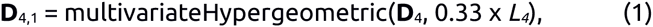

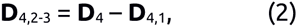

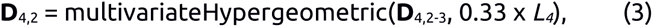

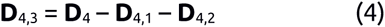

where *L*_4_ is the number of 4-fold sites and **D**_4_ is the total number of 4-fold divergent sites. We divided **D**_4,1_, **D**_4,2_ and **D**_4,3_ by ⅓ × *L*_4_ to get K_4,1_, K_4,2_ and K_4,3_, respectively. We used K_4,1_ to rank genes and assign genes to bins, we then used K_4,2_ to estimate the rate of adaptive non-synonymous substitution (K_a+_) and K_4,3_ as an estimate of the mutation rate.

To test if genes with high mutation rates have lost more adaptive amino acid substitutions than genes with low mutations rates due to HRi, we have categorized genes into low and high mutations groups after splitting K_4_ into two statistically independent variables; K_4,1_ was used to rank genes and assign genes to bins, and K_4,2_ was used to estimate K_a+_. Again, this was done by generating a random multivariate hypergeometric variable as follows:

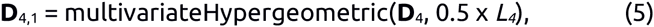

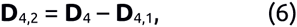

### Estimating the number of substitutions lost to HRi

To estimate how many adaptive substitutions are lost to HRi we proceeded as follows. Let *K*_*a*+(*i*)_, *L*_*a(i)*_ and *RR*_(*i*)_ be the estimated rate of *K*_*a*+_, the total number of 0-fold sites and the average rate of recombination for the *ith* group of genes (grouped by recombination rate). We fit a LOESS curve to the relationship between *K*_*a*+_ and the rate of recombination. Let the estimated value of *K*_*a*+_ for the *ith* group of genes from the LOESS curve be *K*_*a*+(*i*)’_ – this can be thought of as the predicted mean rate of adaptive non-synonymous substitution for genes of the observed recombination rate. We took the average *K*_*a*+_ for genes with rates of recombination above 2 cM/MB as our estimate of the rate of adaptive non-synonymous substitution without HRi – let this be *K*_*a*+, *no_HRi*_. The expected total number of adaptive non-synonymous without any HRi is therefore:

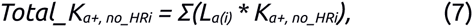

and the number lost to HRi is:

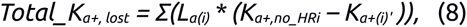

for groups of genes with a rate of recombination < 2 cM/MB. Because the mean rate of adaptive non-synonymous substitution from the LOESS curve can be negative we repeated the analysis setting any value of *K*_*a*+(*i*)’_ to zero if it was less than zero. Finally the proportion of substitutions lost to HRi:

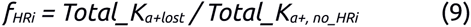

### Confidence Intervals and Bootstraps

To calculate the 95% confidence intervals (CIs) for the proportion of adaptive amino acid substitutions lost due to interference (*f_HRi_*), we bootstrapped 1,000 times the data by gene. We split each 1,000 random data sets into 45 recombination bins (containing 136 genes each) and reestimated K_a+_ for each bin independently using the DFE-alpha software (Eyre-Walker and Keightley 2009, see above). For each random data set, we fitted a LOESS curve to the relationship between K_a+_ and the rate of recombination and reestimated the proportion of substitutions lost to HRi, *f_HRi_* (see above).

For testing if genes undergoing high mutation rates have lost more adaptive substitutions than genes under low mutation rates we bootstrapped the data set by gene 1,000 times. For each bootstrap dataset we split gene’s K_4_ estimates into two variables; K_4,1_ and K_4,2_ sampling from an hypergeometric distribution (see the details of this sampling above). For each bootstrap replicate we: (i) took the 25% of genes with the highest (and lowest) K_4,1_ to define the high (and low) mutation group, (ii) divided each mutation group into 11 recombination rate bins (with 136 genes each) and (iii) estimated K_a+_ using K_4,2_ for each recombination – mutation group. The distribution of *f_HRi_* for each mutation group was obtained by applying expression (9) to each bootstrap replicate. To test if *f_HRi_* differs significantly among mutation groups we estimated the statistic *Z* with the following expression:

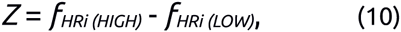

where *f*_*HRi (HIGH)*_ is the proportion of lost substitutions for genes with high mutation rates and *f*_*HRi (LOW)*_ is the proportion of lost substitutions for genes with low mutation rates. Finally, the proportion of the *Z* distribution below zero is the one-tailed p-value for the observed differences between high and low mutation groups.

### Statistical Analyses

All statistical analyses were performed using the R statistical package (R Core Team 2013). ANCOVAs and multiple linear regressions were carried out calling the R function “lm” (from the R package “base”). We calculated Spearman rank correlations (*ρ*) using the basic R function “cor.test” (from the R package “base”). To compare the linear and curvilinear model fit we used the R function “nls” (from the R package “stats”). The random hypergeometric variable was obtained through the R function “rhyper” (from the R package “stats”). LOESS regression was run using the R package “stats” after setting the smoothness parameter “span” from the default 0.75 value to 1. Increasing the “span” parameter decreases the smoothness of the fitted curve making the regressions more robust (or less noisy) across bootstrap replicates. All these R scripts are available upon request.

